# Noise-induced symmetry breaking in a network of excitable ecological systems

**DOI:** 10.1101/2022.08.20.504626

**Authors:** Arzoo Narang, Tanmoy Banerjee, Partha Sharathi Dutta

## Abstract

Noise-induced symmetry breaking has barely been unveiled on the ecological grounds, though its occurrence may elucidate mechanisms responsible for maintaining biodiversity and ecosystem stability. Here, for a network of excitable consumer-resource systems, we show that the interplay of network structure and noise intensity manifests a transition from homogeneous steady state to inhomogeneous steady states, resulting in noise-induced symmetry breaking. On further increasing the noise intensity, there exist asynchronous oscillations, leading to heterogeneity crucial for maintaining a system’s adaptive capacity. The observed collective dynamics can be understood analytically in the framework of linear stability analysis of the corresponding deterministic system.

---

A principle finding in theoretical ecology suggests that even a simple population model can manifest a range of dynamical scenarios, from stable equilibria to cyclic oscillations, through to chaos [1]. Oscillatory dynamics has a long history, as summarized in diverse fields [2–4]. Elucidating various mechanisms behind these oscillations is a major challenge and has been of persistent interest in ecology [5]. As discussed in [6], endogenous causes are the plausible explanation for the generation of population cycles. Another well studied nonlinear phenomenon observed in dynamical systems is the *excitability* [7, 8]. Excitable dynamics are observed in a wide range of natural systems, which under strong perturbations can evoke large-amplitude fluctuations, before relaxing to a rest state [9]. This large amplitude transient fluctuations can sometimes turn into sustained oscillations due to stochastic perturbations, often called noise-induced oscillations [10, 11]. Here, by considering an ecological network of excitable systems, we address the following questions: Do noise-induced oscillations always directly transit from a steady state? Can other intermediate collective dynamics exist while the system shifts from a steady state to noise-induced oscillations?

Stochasticity or noise is ubiquitous in ecosystems. An extensive literature on noise, in recent years, has examined that noise can lead to many novel phenomena, from population cycles to coexistence [12, 13]. The persuasive role of noise on the dynamics of excitable systems is observed in many disciplines, including noise-induced oscillations [10, 14], phenomenon of coherence resonance [15, 16], occurrence of chimera states [17], and noise-enhanced synchronization in coupled excitable systems [18, 19]. An intriguing phenomenon that has received less attention is noise-induced symmetry breaking (NISB). NISB affirms reduced symmetric configuration in the presence of noise, even though the underlying deterministic processes are symmetric, thus resulting in the occurrence of multiple stable states. There is limited research on NISB [20, 21], and so far, its ecological facet remains to be studied.

Multiple stable solutions make it possible for populations in distinct patches/nodes to settle into different steady states. Therefore, minimizing the extinction risk and increasing the stability of spatial population through rescue effect [22]. The idea of spatial ecosystem functioning and species interactions goes hand in hand. Spatially separated populations, which through dispersal may synchronize, are considered necessary to understand population cycles [23]. Researchers find that large systems of interacting oscillators have promising applications in various fluctuating systems [9, 24]. Further, species dispersal network structure is believed to influence the ecological dynamics strongly, as explored by recent studies [25, 26]. Following those lines of thought, here we report that an interplay of network structure and noise intensity results in a transition from homogeneous steady state to inhomogeneous steady states via NISB; before turning into noise-induced asynchronous oscillations. The results are explained numerically with the help of time series, spatiotemporal plots, and phase diagrams. Further, we show that the network model’s linear stability analysis can help explain the observed dynamics.

We consider an ecological network with *N*-patches inhabiting resource-consumer [27] systems. The consumers in each patch are connected with other patches via a diffusive coupling, where the connectivity pattern varies from local to global. There is an additive Gaussian white noise source *ξ*(*t*) that affects the consumer abundance. The network model under stochastic perturbations is read as:

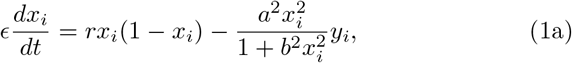

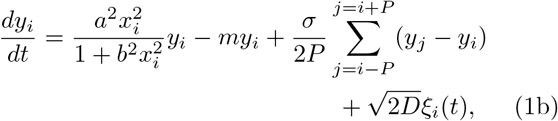

where *x_i_*, and *y_i_*, respectively, determine resource and consumer abundance, *i*(= 1, 2,…, *N*) denotes the patch index, all indices are modulo *N*. The parameter *ϵ* > 0 is responsible for a timescale separation between a fast resource population and a slow consumer population. The resource follows the logistic growth with an intrinsic growth rate *r*, and the interaction of resource and consumer is characterized by Holling’s type-III grazing with parameters *a* and *b*. *m* is the natural mortality of the consumer. The parameter *a* describes the excitability threshold of the isolated system; in particular, it determines whether the system is in the excitable (*a* > 8.975) or the oscillatory (*a* ∈ [7.345, 8.975]) regime. Here, we focus on the dynamics of the resource-consumer population in the excitable regime (*a* = 9). The model assumes the movement of the consumer population between the patches, where the interaction is governed by the coupling strength *σ* and the parameter *P* controls the coupling range *p* = *P/N*, where 1 ≤ *P* ≤ (*N* – 1)/2 for an odd number of patches. Increasing the value of *P* from 1 to (*N* – 1)/2 varies the network topology from local to global via non-local. Further, 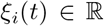 is the normalised Gaussian white noise that perturbs the consumer population in each patch *i*, i.e., 〈*ξ_i_*(*t*)〉 = 0 and 〈*ξ_i_*(*t*)*ξ_j_*(*t*′)〉 = *δ_ij_δ*(*t* – *t*′), ∀*i,j*, and *D* is the noise intensity. In the excitable region, the isolated system rests in a stable steady state in the absence of noise [see Fig. 1(a)]. Inducing stochastic perturbations beyond a threshold value of the noise amplitude drives the population to produce sustained oscillations, as seen in Fig. 1(b).

**FIG. 1.**
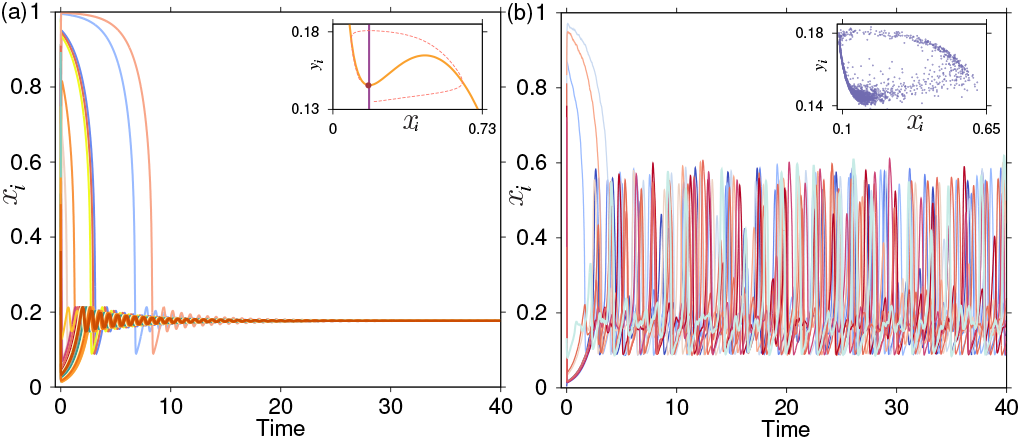
Time series of the resource population (*x*) for the model (1) with, *σ* = 0 and *a* = 9 (the system is in the excitable region): (a) For zero noise intensity (*D* = 0) the system settles into a steady state; inset: nullclines of the resource (*x*) and the consumer (*y*). (b) For a non-zero noise intensity (*D* = 0.00005) there exists noise-induced oscillations; inset: stochastic cyclic attractor. Other model parameters are: *r* = 1, *b* = 7, *m* =1, *ϵ* = 0.01, and *N* = 101.

The interplay of node dynamics, network topology, and noise introduced in the model (1), gives rise to distinct dynamical regimes [28]. Depending upon noise intensity *D*, we demonstrate four distinct space-time patterns in Fig. 2 (*p* = 0.08). In the absence of noise (*D* = 0) or for a low noise intensity, all the nodes rest in the steady state, thus giving a homogeneous steady state solution [Fig. 2(a)]. Now, in the presence of a weak noise strength, the system breaks into two sub-populations having two distinct noise-induced inhomogeneous steady states. The scenario is shown in Fig. 2(b) for *D* = 0.00001. The time series in the right panel of Fig. 2(b) shows two distinct branches of resource densities. Interestingly, the lower branch coincides with the deterministic steady state. However, the presence of noise gives birth to an additional steady state, i.e., the upper branch. The spatiotemporal plot in the left panel of Fig. 2(b) depicts that the upper branch randomly appears in the background of the steady state of the lower branch. The presence of two steady states breaks the spatial symmetry of the system, and we get a new stationary state governed by noise-induced symmetry breaking. We find that the network exhibits stochastic switching between the two branches for an intermediate noise intensity. Fig. 2(c) demonstrates this scenario for *D* = 0.000033. For a large noise intensity (*D* = 0.005), inhomogeneous steady states no longer exist; rather, stochastically spiking incoherent oscillations take place [see Fig. 2(d)].

**FIG. 2.**
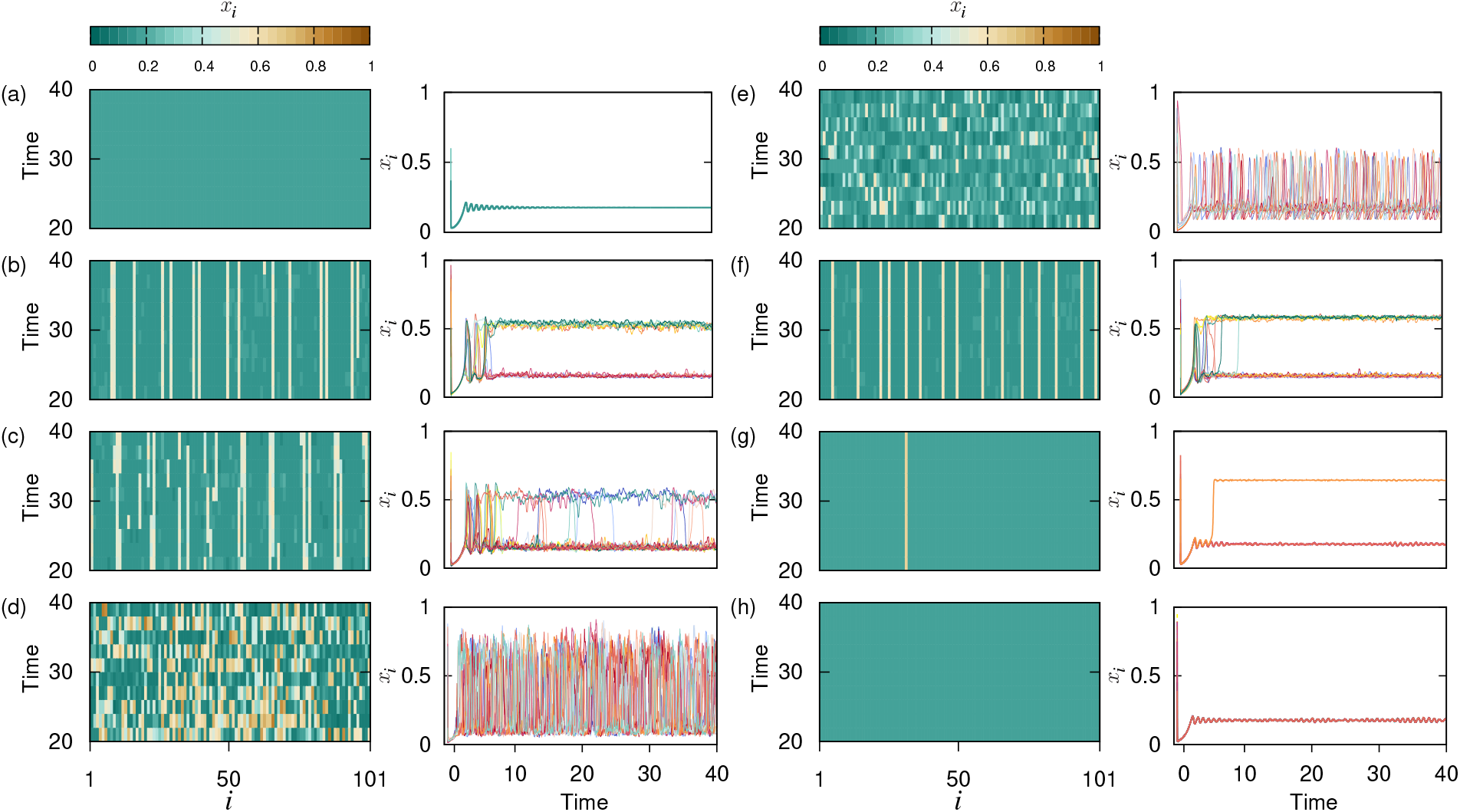
(a)-(d) Space-time (left column) and time series plots (right column) of resource *x_i_* for *P* = 8, *σ* = 0.1 with varying noise intensities: (a) *D* = 0; steady state, (b) *D* = 0.00001; symmetry breaking, (c) *D* = 0.000033; stochastic switching between two resource densities, and (d)*D* = 0.005; asynchronous oscillations. (e)-(h) Space-time (left column) and time series (right column) plots for *D* = 0.00001, *σ* = 0.6 with varying coupling range *p*: (e)*p* = 0.01 (local coupling); asynchronous oscillations, (f) *p* = 0.04; symmetry breaking, (g) *p* = 0.25; symmetry breaking with most nodes settling at the lower branch, and (h): *p* = 0.5 (global coupling); steady state. Other model parameters are *r* = 1, *a* = 9,*b* = 7, *m* = 1, *ϵ* = 0.01, and *N* = 101.

Moreover, to provide an insight of the effects of network topology on the dynamics of the system, we fix the noise intensity *D* = 0.00001, coupling strength *σ* = 0.6 and vary the connectivity from local (*p* = 0.01) to global (*p* = 0.5) via non-local (*p* = 0.03, *p* = 0.25) coupling [see Figs. 2(e)-2(h)]. We find notable differences between dynamics with the changing network topology. Increasing the coupling range of the network not only reduces the number of solutions but also changes the dynamics from oscillatory to steady state. As observed from Fig. 2(e), local coupling favors oscillating populations with spatial incoherence, whereas increasing the coupling range shows a transition from oscillating populations [Fig. 2(e)] to inhomogeneous steady states [Figs. 2(f)-2(g)], through to homogeneous solutions [Fig. 2(h)]. Interestingly, in the presence of non-local coupling, as depicted in the spacetime plot in Fig. 2(f), oscillators randomly rest at the lower or the upper branch; however, with the increasing network connectivity to global coupling, all the oscillators settle at the lower branch [Fig. 2(h)], which also incites towards homogeneous population densities. Thus, for a fixed noise intensity *D* and coupling strength *σ*, the network model (1) experiences NISB while moving from global to local coupling.

The observed transition from oscillatory dynamics to inhomogeneous states impels us to investigate our system’s qualitative behavior in these respective regimes. We analyze the features of the oscillatory region through phase portrait in Figs. 3(a)-3(b) for noise intensity *D* = 0.005. We observe that the density of phase points of stochastic limit cycle attractor for one oscillator [see Fig. 3(a)] and correspondingly for all the oscillators [see Fig. 3(b)] is larger around two population densities, i.e., *x_i_* = 0.1 and *x_i_* = 0.55. To further elaborate on the phenomenon of symmetry breaking observed in Fig. 2, we calculate the center of mass defined as [29]:

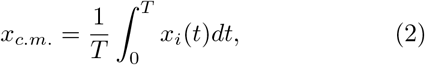

where *x_i_* is the resource density in each *i*-th patch, and *T* is taken sufficiently large. Resource population settles exactly into two branches as is characterized by two distinct values of center of mass [Fig. 3(c)], where for one part of the population *x_c.m_*. ≈ 0.15 and for the other subpopulation *x_c.m_*. takes the value around 0.5, therefore exhibiting two nonuniform states. A relevant observation illustrated in the phase space [Fig. 3(d)] suggests an underlying mechanism for symmetry breaking, specifically NISB, clearly indicating the coexistence of two steady states in the network. We observe that wide distributions show up for two density values (≈ 0.15 and ≈ 0.5), thus settling the system solely around these two states.

**FIG. 3.**
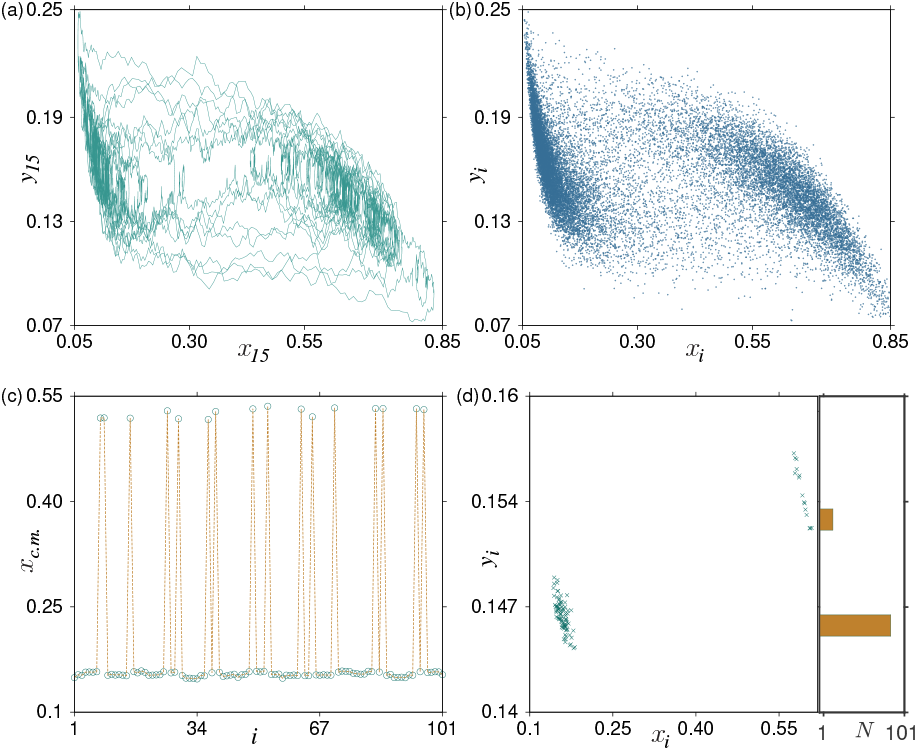
Phase portraits exhibiting the long stay of attractor in the two domains around *x_i_* = 0.1 and *x_i_* = 0.55 for (a) one oscillator (15th node), and (b) all the oscillators, at noise intensity *D* = 0.005. (c) Center of mass (*x_c.m_*.), and (d) phase portrait at a particular time along with density distribution of *y_i_* corresponding to NISB for non-local coupling (*P* = 8) at noise intensity *D* = 0.00001. The histogram in the right panel of (d) represents the number of oscillators in either of the two states. Other model parameters are *r* = 1, *a* = 9, *b* = 7, *m* = 1, *ϵ* = 0.01, *σ* = 0.1, and *N* = 101.

To gain a comprehensive view of the spatiotemporal dynamics in the network, we compute phase diagrams in the (*p, D*) and (*p, σ*) parameter planes [see Figs. 4(a) and 4(b), respectively]. Keeping the value of coupling strength *σ* fixed to 0.1, we vary *p* and *D* in Fig. 4(a). For stronger noise intensity *D*, in the entire range of *p*, the system resides in the asynchronous oscillatory regime. For weaker values of *D*, we observe either asynchronous oscillations or NISB, along with a region of stochastic switching, depending upon the coupling range *p*. NISB occurs for a certain threshold value of *p* > 0.07, i.e., when the coupling is non-local with around 8 nodes connected and persists up to *p* = 0.5 (globally coupled). However, local coupling (*p* = 0.01) and less connected nodes (up to 6) maintain the oscillatory behavior. Moreover, a narrow zone of stochastic switching is observed between these two regimes. In Fig. 4(b), we explore the interplay of *p* and *σ*, with value of *D* being fixed to 0.00001. The oscillatory region observed for a large number of connected nodes *p* = 0.21 narrows down to *p* = 0.02 with increasing coupling value, clearly determining the persisting oscillatory pattern for local coupling in the whole range of *σ*. Moreover, for a large value of *σ* (≈ 0.53), a transition from oscillations to NISB via stochastic switching takes place for a lower coupling range. In contrast, with decreasing coupling strength, more connected nodes are required for the transition. In the direction of global coupling, beyond a threshold value of *σ*, the system traverses a synchronous steady state (HSS).

**FIG. 4.**
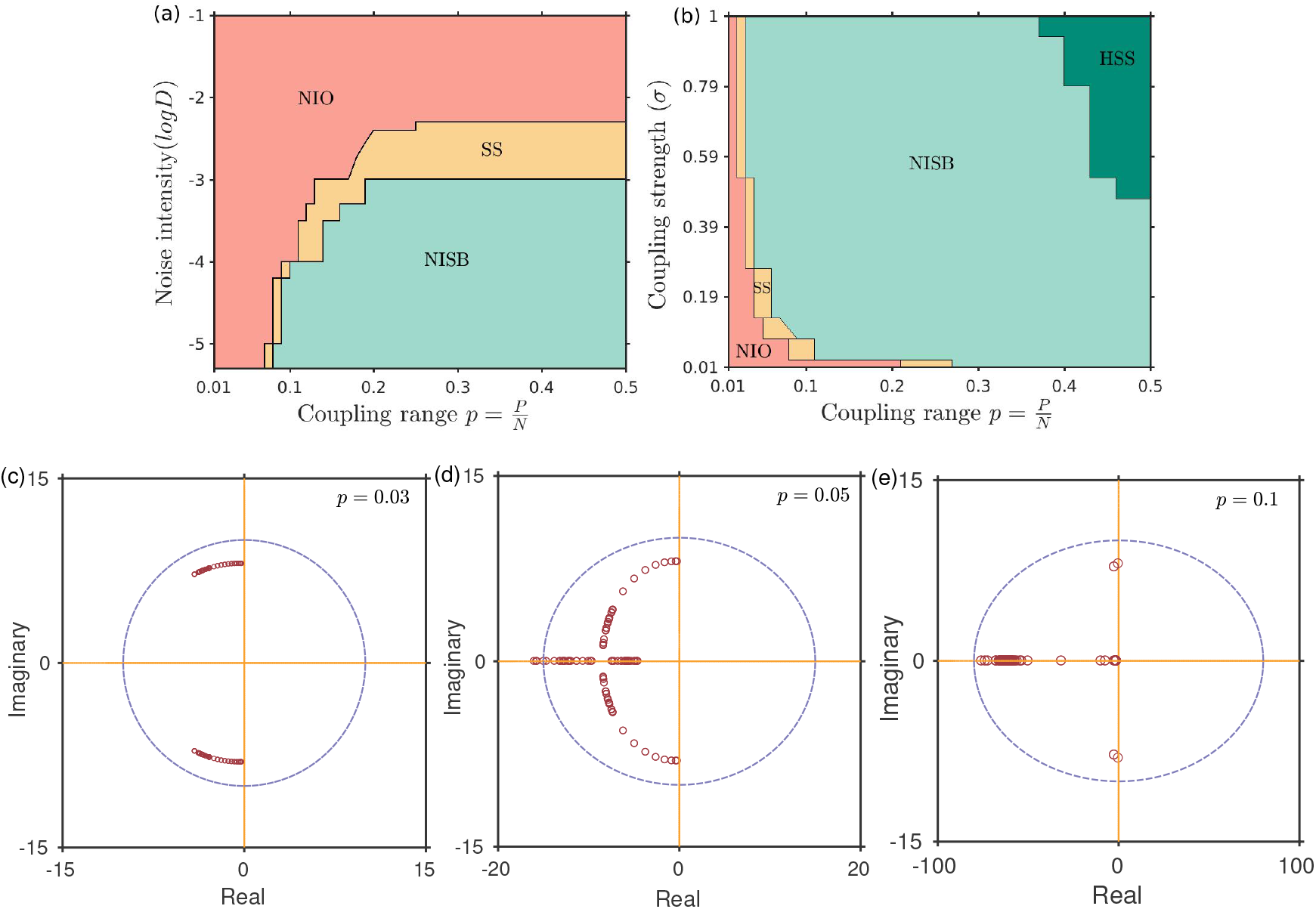
Phase diagrams in the (a) (*p, D*) plane for *σ* = 0.1, and (b) (*p*, σ) plane for *D* = 0.00001, where NIO: noise-induced oscillations, SS: Stochastic switching, NISB: noise-induced symmetry breaking, and HSS: Homogeneous steady state. (c)-(e) Distribution of the eigenvalues for different values of *p* for *σ* = 0.6 and *D* = 0. Other model parameters are same as in Fig. 2.

The noise intensity *D* considered for our analysis is in the order of 10^-5^ [see Fig. 2]. Thus, insinuating the perceived outcomes in the stochastic system might be the effect of the deterministic skeleton. Therefore, considering the deterministic framework, i.e., *D* = 0 in (1), we carry the linear stability analysis. We calculate eigenvalues of the linearized system using the equilibrium points based upon the changing network topology [30]. Let 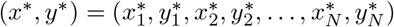 be a non-trivial equilibrium point of the system (1) when *D* = 0, and it varies with the coupling range (*p*). Linearization of the deterministic system in the neighborhood of the equilibrium point (*x*\*, *y*\*) yields the following block-structured matrix:

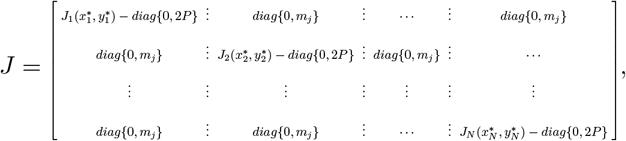

where *J_i_* is the Jacobian of the isolated *i*-th patch at an equilibrium point 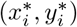 (*i* = 1,2,…, *N*) given as:

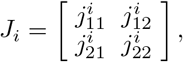

with 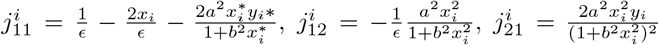 and 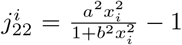. Further *m_j_* = 1 if *i*-th and *j*-th nodes are connected, and *m_j_* = 0 otherwise.

The coupling strength *σ* = 0.6 and noise intensity *D* = 0.00001 manifest four distinct regimes based upon the value of *p*, as can be seen in Fig. 4(b). The transition from oscillatory to NISB via SS occurs for *p* = 0.05, further traversing HSS around *p* = 0.41. We here intend to investigate whether the node dynamics and network structure determine the dominant pattern of NISB in the limiting value of *D* tending to 0. Figs. 4(c)-4(e) show the distribution of eigenvalues with varying coupling range *p*. We observe complex conjugate eigenvalues (*λ_i_*) for *p* ranging from 0.01 to 0.04 as shown for *p* = 0.03 in Fig. 4(c). It is ascertained from Fig. 4(c) that the fixed point obtained at *D* = 0 for *p* = 0.03 is a stable spiral since *Real*(*λ_i_*) < 0 ∀*i*. However, as analyzed from Fig. 4(b) for the same coupling range (*p* = 0.03), the presence of noise in the system results in the occurrence of oscillations or SS. A recent study [31] demonstrated the impact of additive noise on networks, tuning their spectral properties. The work showed that increasing noise intensity could coerce the eigenvalues to cross the imaginary axis. Therefore, we infer that the transition from a steady state (*D* = 0) to an oscillatory state or a region of SS (*D* ≠ 0) in our work is due to the impact of noise on the eigenvalue spectrum that resulted in destabilizing the equilibrium point by crossing the imaginary axis, and hence the occurrence of Hopf bifurcation. Moving further to *p* = 0.05, we observe the emergence of a few real eigenvalues [see Fig. 4(d)], which also eventually tend towards 0 with increasing coupling range (*p* = 0.1), as observed from Fig. 4(e). We also notice that the number of real eigenvalues increases in a passage of global coupling. Thus, we infer that the presence of noise in the network (1) can lead the real eigenvalues to cross the origin, henceforth inducing NISB via a pitchfork bifurcation.

In conclusion, we have shown that an ecological network of identical excitable systems can be driven out of the resting state leading to different collective dynamics, including regimes of heterogeneous steady states and asynchronous oscillatory states, mediated by noise intensity and network topology. For non-local coupling and adequately tuning the noise intensity, we achieve an oscillatory regime due to the resonance effect [11] or a region of inhomogeneous steady states (NISB). Further, by keeping the noise intensity fixed while changing the coupling range from local to global, we perceive a transition from an oscillatory domain to a region HSS through NISB. Although the oscillatory and inhomogeneous steady states promote species’ sustainability, the emergence of HSS while traversing towards global coupling inhibits biodiversity [26]. Our results are robust across a large region in the parameter space, as demonstrated via phase diagrams. Therefore, our findings could be important for understanding the mechanisms responsible for upholding biodiversity and ecosystem stability.

## ACKNOWLEDGMENTS

P.S.D. acknowledges financial support from the Science & Engineering Research Board (SERB), Government of India (Grant number: MTR/2021/000148).

